# Evolution of multicellularity and reproductive strategies in yellow-green algae (Xanthophyceae, Heterokontophyta)

**DOI:** 10.64898/2026.08.06.743135

**Authors:** Seok-Wan Choi, Paul A. Broady, Phil M. Novis, Robert A. Andersen, Hwan Su Yoon

## Abstract

The evolution of multicellularity has long been linked to reproductive strategies. A long-standing debate concerns whether multicellular organisms are primarily stabilized by small single-cell propagules that minimize genetic heterogeneity or by larger multicellular and multinucleate propagules that may improve developmental success and survival of individuals. the Xanthophyceae provides an excellent model for investigating these questions, exhibiting transitions between unicellular to multicellular filamentous and coenocytic forms together with diverse reproductive modes, including single-cell zoospores and autospores, and multinucleate monospores and akinetes. However, a robust phylogenetic framework and systematic analyses of character evolution have remained lacking in this lineage. Here, we present a phylogenomic framework based on a nuclear dataset of 680 genes from 18 species, including 17 newly generated transcriptomes. Nuclear phylogenies robustly resolve all sampled inter-ordinal and inter-familial relationships with full concordance between concatenation and coalescent analyses, while plastid (141 genes) and mitochondrial (31 genes) datasets from 33 species recover identical topologies. Based on these results, we establish one new order (Pseudopleurochloridales), emend one order (Heterococcales), and propose five new families. Ancestral character reconstruction indicates at least four independent transitions from unicellular ancestors to simple multicellularity. Bayesian analyses of multicellularity and reproductive characters show that these transitions were consistently accompanied by shifts from multiple autospore-type propagules toward single monospore- and akinete-type propagules, whereas reversions to unicellularity were associated with the reappearance of autospore-based reproduction. These results provide a phylogenomic framework for understanding multicellular evolution in Xanthophyceae and shed light on the relationship between reproductive modes and the emergence of simple multicellularity.

## 1. Introduction

Multicellularity represents a major evolutionary transition in which previously free-living unicellular organisms gave rise to integrated multicellular organisms (Smith and Szathmáry 1997; Grosberg and Strathmann 2007). A central challenge in this transition is the alignment of cell-level and organism-level fitness (Michod and Roze 1997; Michod 2005). If genetic heterogeneity and competition among cells are not constrained, multicellular organization can be destabilized by within-organism conflict. Reproductive mode is therefore expected to play a key role in the stabilization of multicellularity, because life cycle structure determines how genetic heterogeneity is transmitted, reduced, or reorganized between generations (Haccou and Schneider 2004).

A long-standing view emphasizes the importance of a unicellular bottleneck (Grosberg and Strathmann 1998). Development from a single cell can reduce within-organism genetic heterogeneity, increase relatedness among cells, and limit the success of selfish cell lineages. Consistent with this view, theoretical models have shown that larger multicellular propagules may increase mutation load when the cells that initiate offspring are genetically heterogeneous or distantly related (Kondrashov 1994). However, unicellular propagules are not the only possible reproductive solution. Multicellular organisms may also reproduce through vegetative fragments, multicellular, or multinucleate propagules. These larger propagules may be advantageous when increased offspring size improves establishment, growth, developmental success, or survival. Thus, reproductive evolution during the origin of multicellularity can be viewed as a trade-off between the genetic benefits of small single-cell propagules and the developmental or ecological advantages of larger propagules. Which strategy is favored may depend on the strength of selfish mutations, the benefits of offspring size, organismal growth constraints, and the extent of cell- or nuclear-level selection within an individual (Otto and Orive 1995; Roze and Michod 2001).

Despite extensive theoretical work, empirical tests of the relationship between multicellularity and reproductive mode remain limited, especially outside animals, green algae, and a few experimentally tractable model systems (Ratcliff et al. 2013; Brunet and King 2017). Protist lineages are particularly important in this context because many contain living forms that span the transition from unicellular organization to simple multicellularity. Such lineages can provide comparative systems for testing whether changes in body organization are repeatedly associated with shifts in reproductive strategy.

Xanthophyceae, or yellow-green algae, provides a useful but understudied system for addressing this question. They belong to the Heterokontophyta (photosynthetic stramenopiles; Guiry et al. 2023) lineage that originated through secondary endosymbiosis of a red algal plastid more than 800 million years ago (Yoon et al. 2004; Parfrey et al. 2011; Strassert et al. 2021; Choi et al. 2024). Xanthophyceae are closely related to Phaeophyceae (brown algae; Yang et al. (2012)), but differ markedly in ecology and body-plan evolution. Whereas brown algae evolved complex multicellularity mainly in marine environments (Denoeud et al. 2024), Xanthophyceae are predominantly freshwater and terrestrial (only ∼5% marine or brackish; 34 spp.). and include a broad range of organizational forms, from unicellular coccoid species to filamentous and coenocytic thalli (Fig. 1a; Maistro et al. (2017)). This diversity makes Xanthophyceae one of the few photosynthetic stramenopile lineages in which repeated transitions between unicellular and simple multicellular organization can be examined within a comparative phylogenetic framework.

**Figure 1.**
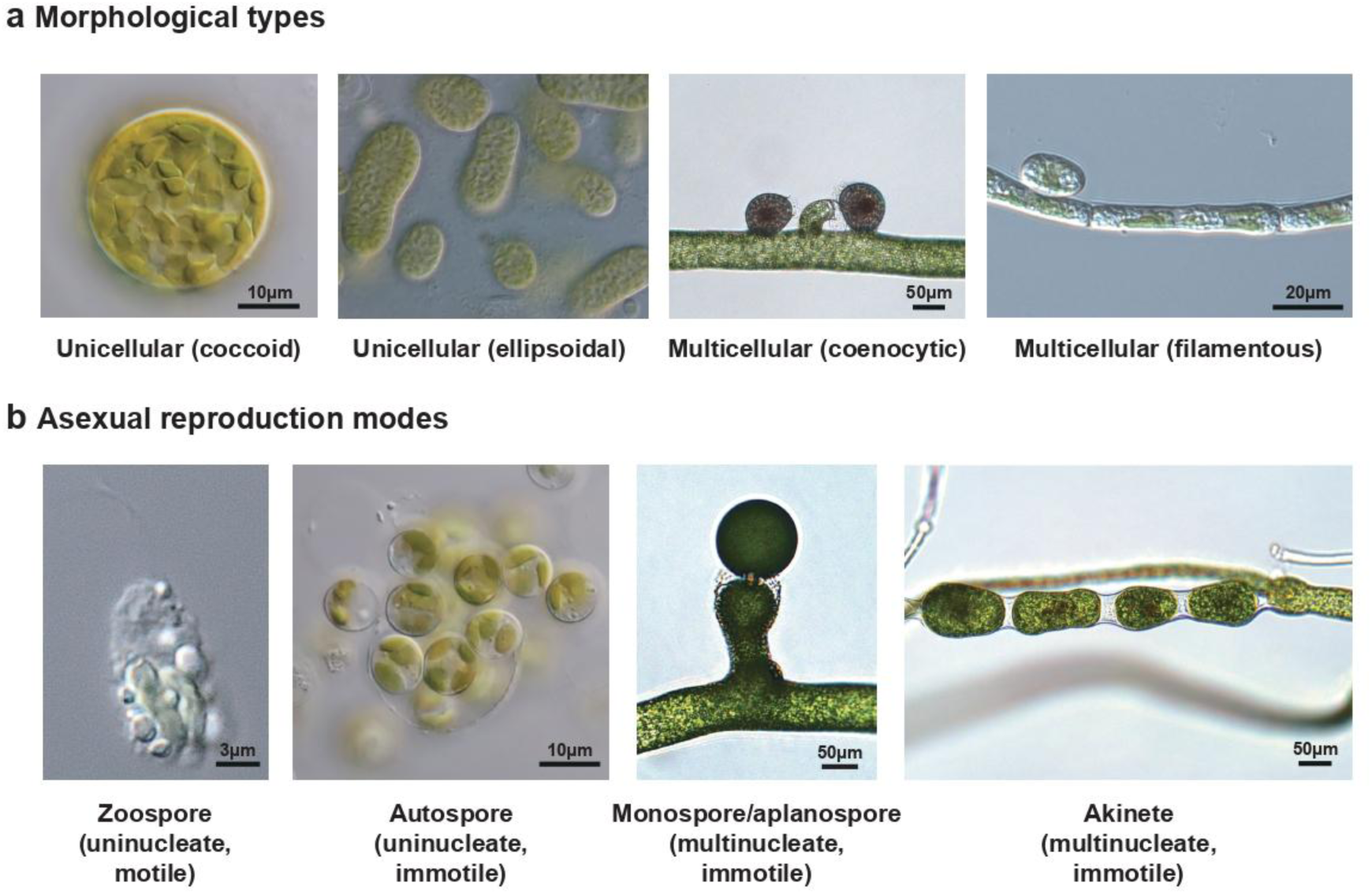
Diversity of cell organization and asexual reproductive modes in Xanthophyceae and related heterokontophytes. (a) Representative unicellular and multicellular morphologies (from left to right: *Botrydiopsis pyrenoidosa*, *Excentrochloris constricta*, *Vaucheria bursata*, and *Bumilleria sicula*). (b) Representative asexual reproductive structures and propagule types (from left to right: the two leftmost images, *Botrydiopsis pyrenoidosa*; *Vaucheria canalicularis*; and *Asterosiphon dichotomus*).

Xanthophyceae and related lineages are also notable for their diversity of asexual reproductive modes. Many taxa produce motile zoospores, while non-motile propagules include autospores, monospores (single-cell aplanospores defined herein; refer to Appendix for definitions), and akinetes (Fig. 1b). These reproductive structures differ in cell number, nuclear condition, motility, and developmental processes. In particular, the contrast between multiple, small, mononucleate autospore-type propagules and single, larger, often multinucleate monospore- or akinete-type propagules provides an opportunity to examine whether the emergence of simple multicellularity is associated with shifts in propagule strategy. Thus, Xanthophyceae offer an important system for extending theories of propagule-size evolution and unicellular bottlenecks to a natural protist lineage.

Historically, the classification of Xanthophyceae was based mainly on vegetative morphology. Traditional systems recognized orders corresponding to flagellate, amoeboid, palmelloid, coccoid, filamentous, and siphonous/coenocytic forms (Pascher 1939; Ettl 1978; Silva 1979). However, molecular studies based on multigene datasets (18S rRNA, *rbc*L, *psa*A) showed that many morphology-based orders, families, and genera are para- or polyphyletic, reflecting extensive morphological convergence and plasticity (Potter et al. 1997; Bailey and Andersen 1998; Negrisolo et al. 2004; Maistro et al. 2009). Although these studies greatly improved our understanding of xanthophyte diversity, sparse gene and taxon sampling left deep relationships unresolved and limited the use of the group for robust evolutionary reconstruction.

A robust phylogenetic framework is an essential prerequisite for testing hypotheses on the evolution of multicellularity and reproductive strategies. Accordingly, we first establish the first genome-scale phylogeny of Xanthophyceae before reconstructing character evolution. We present a phylogenomic framework for Xanthophyceae, based on nuclear, plastid, and mitochondrial datasets, including 17 newly generated transcriptomes and 28 newly generated organelle genomes. This framework resolves major inter-ordinal and inter-familial relationships (Fig. 2), providing a basis for higher-level taxonomic revision, and enables systematic reconstruction of character evolution. Using this phylogeny, we test whether transitions to simple multicellularity were associated with shifts in reproductive mode, particularly from multiple autospore-type propagules toward single monospore- and akinete-type propagules. By linking phylogenomics, morphology, and reproductive traits, this study sheds light on the correlated evolution of reproductive strategies and simple multicellularity in a poorly understood photosynthetic protist lineage.

**Figure 2.**
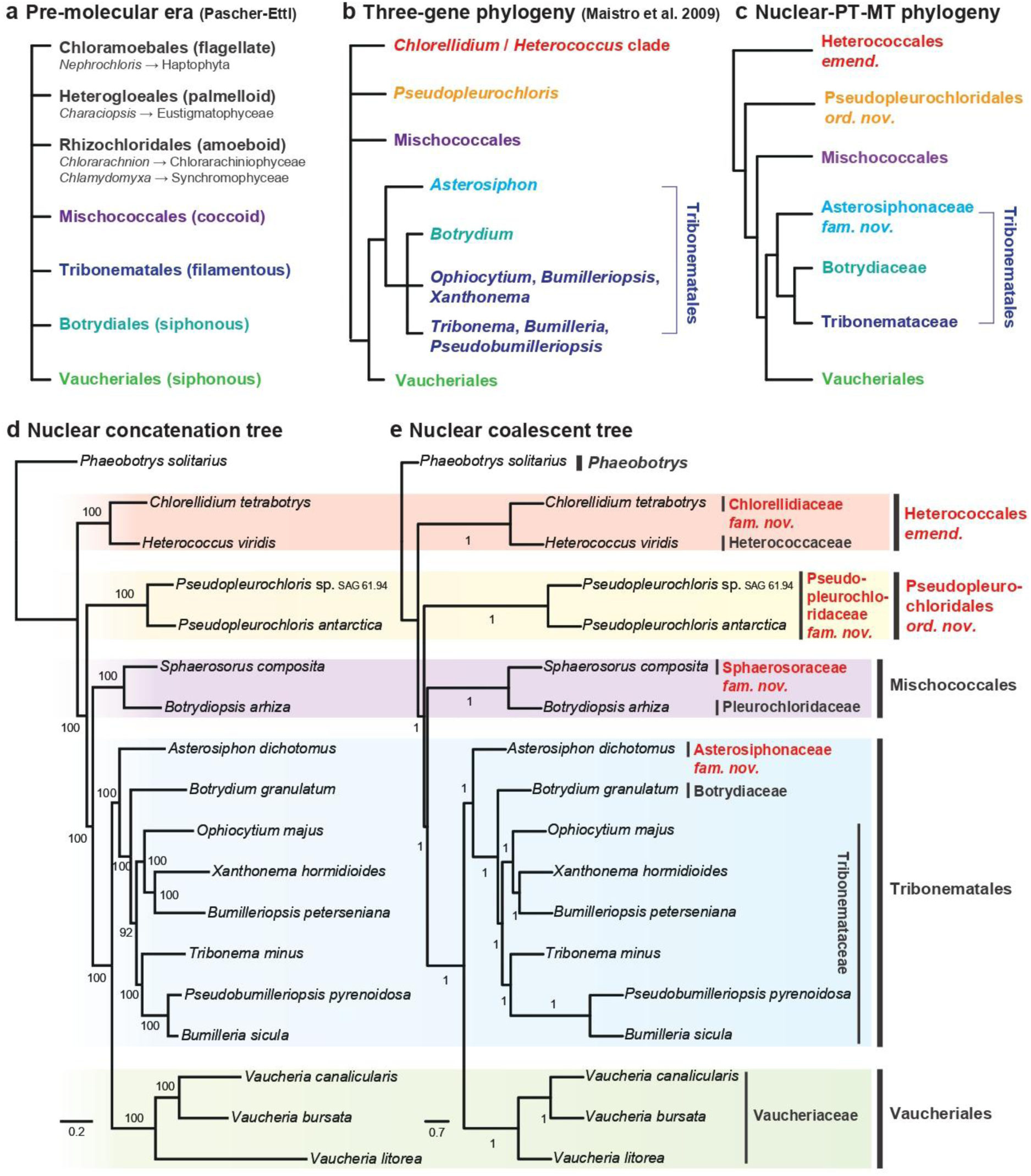
Phylogeny and taxonomy of Xanthophyceae. (a) Traditional taxonomy of Xanthophyceae in the pre-molecular era (after Ettl, 1978). (b) Three-gene phylogeny (SSU rDNA, *rbc*L, *psa*A) from Maistro et al. (2009), with branches showing <70% bootstrap support collapsed into polytomies. (c) Fully resolved phylogeny inferred in this study with updated taxonomic framework. (d) Nuclear concatenation tree and (e) species tree inferred under the coalescent model, both based on 680 nuclear genes. Newly proposed taxonomic revisions are highlighted in red. Ultrafast bootstrap (UFBoot) values and local posterior probabilities are shown on branches in (d) and (e), respectively.

## 2. Materials and methods

### Sampling, culture acquisition, and culturing conditions

A total of 28 strains were included in this phylogenomic and phylotranscriptomic study, comprising 24 xanthophytes together with *Phaeobotrys solitarius*, *Nematochrysis sessilis*, *Phaeothamnion wetherbeei* (Phaeothamniophyceae), and *Choristocarpus tenellus* (Phaeophyceae). 14 strains were obtained from the Culture Collection of Algae at the University of Göttingen, Germany (SAG), three from the Norwegian Culture Collection of Algae (NORCCA), two from the Culture Collection of Algae at the University of Texas at Austin (UTEX), and one from the Kobe University Macroalgal Culture Collection (KU-MACC). Additional strains were derived from private cultures maintained by the authors, including *Botryochloris* sp. (PAB 938) and *Chlorellidium pyrenoidosum* (PAB 785) from P.A.B.; two *Botrydiopsis constricta* strains (NCBC and NCBP) and *Pseudopleurochloris* sp. (LCR-Ant16_1) from P.M.N.; *Nematochrysis sessilis* (A14,626) from R.A.A.; and *Vaucheria bursata* (V-0031) from S.-W.C. and H.S.Y. Detailed information on strain identity, taxonomy, sampling locality, collection dates, and collectors/isolators is provided in Supplementary Table S1.

The marine brown alga *Choristocarpus tenellus* (KU-1152) and xanthophyte *Sphaerosorus compositus* (SAG 53.91), as well as *Nematochrysis sessilis* (A14,626), were cultured in enriched seawater L1 medium (Guillard and Hargraves 1993), with ammonium added to a final concentration of 500 μM for *N. sessilis*. *Botrydiopsis* isolates and *Pseudopleurochloris* sp. collected by P.M.N. were grown in 50% BG-11 and 10% BG-11 media, respectively. All other strains were freshwater taxa grown in DY-V medium adjusted to pH 6.8. Cultures were maintained at 15°C under a 12:12 h light:dark photoperiod.

### DNA/RNA extraction, sequencing, and data acquisition

Genomic DNA was extracted from samples frozen in liquid nitrogen and pulverized using an Automill TK-AM5 frozen crusher (Tokken Inc., Kashiwa, Japan). DNA and RNA were extracted using the DNeasy Plant Mini Kit and RNeasy® Plant Mini Kit (Qiagen, Hilden, Germany), respectively, following the manufacturer’s protocols. DNA extraction methods for the New Zealand *Botrydiopsis* and Antarctic *Pseudopleurochloris* strains were described previously (Novis et al. 2008; Novis et al. 2015). Libraries with an average insert size of approximately 550 bp were prepared using the TruSeq DNA Nano protocol for genomic DNA and the TruSeq Stranded mRNA protocol for RNA sequencing. The NEBNext® Ultra™ II FS DNA Library Prep Kit for Illumina (McKay et al. 2019) was used for DNA library preparation of low-concentration PAB samples. Whole-genome and transcriptome sequencing were carried out on an Illumina NovaSeq X Plus platform (Illumina, San Diego, CA, USA). Library preparation and sequencing were conducted by JSLINK Co. (Seoul, Korea). In addition, raw genomic sequencing data for *Heterococcus protonematoides* (UTEX 2798) and *Tribonema minus* (UTEX B 3156) were retrieved from the NCBI Sequence Read Archive (SRA; accessions SRR14099987 and SRR14114641, respectively). Three genes (18S rRNA, *rbc*L, *psa*A) from 67 xanthophyte and related species were obtained from the NCBI nr database or newly sequenced in this study. Species information and accession numbers are provided in Supplementary Table S2.

### Genome and transcriptome assembly, organelle annotation, and ortholog inference

Illumina genomic reads were assembled using SPAdes v4.0.0 (Bankevich et al. 2012), and transcriptome reads were assembled using rnaSPAdes (Bushmanova et al. 2019) implemented in the same package. Assembled genome contigs were screened for putative organelle-derived sequences using tblastn v2.2.31 (Camacho et al. 2009), and identified contigs were used as seeds for organelle genome assembly with NOVOPlasty v3.7 (Dierckxsens et al. 2017), followed by circularization. Assemblies were polished by mapping raw sequencing reads using Bowtie2 v2.3.5.1 (Langmead and Salzberg 2012) and processing alignments with SAMtools v1.5 (Li et al. 2009). Final organelle genome assemblies were annotated using GeSeq v2.03 (Tillich et al. 2017) and manually curated in Geneious Prime 2019.0.3 (Kearse et al. 2012). Transfer RNA genes were annotated with tRNAscan-SE v2.0.7 (Chan et al. 2021).

Transcriptome assemblies were processed to predict coding regions using TransDecoder v5.5.0. First, candidate open reading frames (ORFs) were identified using TransDecoder.LongOrfs. The resulting ORFs were searched against the UniProt Swiss-Prot database (Consortium (2019); downloaded March 16, 2021) using blastp v2.231, and the hits were used to guide coding sequence prediction with TransDecoder.Predict, which retains biologically supported ORFs based on homology evidence. Redundant sequences were subsequently reduced using CD-HIT-EST v4.7 with a 99% identity threshold (Fu et al. 2012). The resulting nuclear protein sequences were evaluated using BUSCO v5.8.0 with the eukaryota_odb10 database (Simão et al. 2015).

Orthologous groups were inferred using OrthoFinder v3.0.1b1 (Emms and Kelly 2019) based on a core set of six representative species selected for their high BUSCO completeness scores, characterized by high proportions of complete single-copy genes and low levels of gene duplication: *Phaeobotrys solitarius*, *Heterococcus viridis*, *Pseudopleurochloris* sp. SAG 61.94, *Botrydiopsis arhiza*, *Asterosiphon dichotomus*, and *Vaucheria canalicularis*). To expand ortholog coverage across additional taxa, ortholog sequences were propagated using a hidden Markov model (HMM)-based approach. For each orthogroup, multiple sequence alignments were generated with MAFFT v7.526 (Katoh and Standley 2013), and profile HMMs were constructed using hmmbuild (HMMER v3.1b2; Eddy (2011)). These profiles were then searched against proteomes of 12 additional species using hmmsearch, and the top-scoring hits passing an e-value threshold (1e-20) were extracted and incorporated into the orthogroups. Each gene tree was manually inspected to identify potential contamination or misassignment to orthogroups. This procedure resulted in a combined dataset of 18 species.

### Phylogenomic analyses

For reconstruction of the nuclear maximum-likelihood (ML) phylogeny, we used a concatenated dataset of 680 orthologous nuclear proteins from 18 species. Each gene was aligned individually using MAFFT v7.526 (Katoh and Standley 2013) with the --maxiterate 1000 option, and the resulting alignments were trimmed using trimAl (v1.2rev59) with the -gt 0.9 option (Capella-Gutiérrez et al. 2009) and concatenated. The concatenated matrix was partitioned by gene. The best-fitting amino acid substitution models were selected using ModelFinder (Kalyaanamoorthy et al. 2017), and ML phylogenies were inferred with IQ-TREE v2.3.5 (Minh et al. 2020), allowing each partition to have its own branch length (Chernomor et al. 2016). Branch support was assessed using 1,000 ultrafast bootstrap (UFBoot) replicates (Hoang et al. 2018).

For coalescent-based species tree inference, individual gene trees were first reconstructed using IQ-TREE. Branches with less than 10% bootstrap support were collapsed prior to analysis. The resulting gene trees were then used as input for ASTRAL v5.7.8 (Zhang et al. 2018) to estimate the multispecies coalescent tree, with branch support evaluated using local posterior probabilities (Sayyari and Mirarab 2016).

To assess congruence across genomic compartments, phylogenetic analyses were also conducted using plastid (141 genes) and mitochondrial (31 genes) datasets, expanded to 33 species. Organelle phylogenies were reconstructed using the same alignment, trimming and ML inference procedures as described for the nuclear dataset.

For family- and genus-level taxonomic investigations, a taxon-rich phylogeny encompassing available sequences from Xanthophycean and related genera was constructed using three genes (18S rRNA, *rbc*L, and *psa*A). Phylogenetic reconstruction followed the same alignment and ML inference procedures described above, except that the alignments were manually trimmed prior to analysis.

Alternative phylogenetic hypotheses at ordinal and familial levels within Xanthophyceae were evaluated using topology tests implemented in IQ-TREE. Constrained trees representing alternative relationships were compared using resampling estimated log-likelihoods (RELL; 10,000 replicates), including bootstrap proportion (BP, Kishino et al. (1990)), weighted Kishino–Hasegawa (KH; Kishino and Hasegawa (1989)), weighted Shimodaira–Hasegawa (SH; Shimodaira and Hasegawa (1999)), expected likelihood weight (ELW; Strimmer and Rambaut (2002)), and approximately unbiased (AU; Shimodaira (2002)) tests.

### Quantification of gene-tree characteristics and statistical analyses

To quantify variation in phylogenetic signal among loci, we calculated a set of branch-based metrics for each gene tree following the framework of Vankan et al. (2022). We analyzed the gene trees using ETE3 Python toolkit (Huerta-Cepas et al. 2016). For each gene tree, we computed: (1) the coefficient of variation (CoV) of root-to-tip distances, measured from a midpoint-rooted tree, as an indicator of among-lineage rate heterogeneity; (2) tree length, defined as the sum of all branch lengths, representing the overall substitution rate; (3) stemminess, calculated as the ratio of internal to terminal branch lengths, reflecting tree shape; and (4) mean branch support, calculated as the average of support values across internal nodes, representing the consistency of phylogenetic signal. To assess topological congruence, we calculated the normalized Robinson–Foulds (RF) distance (Robinson and Foulds 1981; Penny and Hendy 1985) between each gene tree and the species tree, as well as the mean RF distance between each gene tree and all other gene trees.

To evaluate the impact of gene selection on species tree inference, loci were ranked based on individual metrics. For each dataset, subsets corresponding to the top 20%, 40%, 60%, and 80% of loci were selected. Loci were selected in ascending order for metrics where lower values indicate higher quality phylogenetic signal (CoV) or more proper for resolving ancient, deep divergences (tree length), and in descending order for metrics where higher values are desirable (stemminess and mean branch support). Selected loci were concatenated into supermatrices, and phylogenetic trees were reconstructed using IQ-TREE v2.3.5 (Minh, Schmidt, et al. 2020).

To examine relationships among gene-tree metrics, we performed correlation analyses using Spearman’s rank correlation coefficients implemented in R. Pairwise correlations were calculated among all variables, including CoV, tree length, stemminess, mean branch support, and RF-based distances. Results were visualized using correlation heatmaps and pairwise scatterplots to identify potential dependencies or redundancies among explanatory variables.

To evaluate the relative contributions of gene-tree characteristics to phylogenetic accuracy, we performed multiple linear regression analyses in R. The normalized RF distance between gene trees and the species tree was used as the response variable, and gene-tree metrics (CoV, tree length, stemminess, and mean branch support) were included as explanatory variables. Model significance was assessed using F-statistics, and the strength and direction of individual predictors were evaluated using regression coefficients and associated *p*-values. This approach allowed us to determine which aspects of branch-length variation and phylogenetic signal most strongly influence gene-tree concordance with the species tree.

### Ancestral character estimation and transition analyses

Ancestral character state estimation of morphological traits was conducted using the maximum-likelihood (ML) phylogram inferred from a taxon-rich three-gene dataset (18S rRNA, *rbcL*, and *psaA*), with the backbone topology constrained to the well-supported chloroplast genome phylogeny to maximize taxon sampling. Species were classified into discrete morphological states representing unicellular and multicellular forms based on previously documented life cycle characteristics (see Supplementary Table S3 and Appendix for character definitions and descriptions). Analyses were performed in R using the package phytools v2.5.2 (Revell 2012). Discrete character evolution models, including equal-rates (ER) and all-rates-different (ARD), were fitted using the function fitMk, and model fit was compared using Akaike Information Criterion (AIC). Stochastic character mapping was then performed under each model using sinmap with 1,000 simulations, and posterior probabilities of ancestral states at each node were summarized from the posterior distribution. The best-fitting model was selected based on AIC and used for downstream visualization and interpretation.

To further quantify transition dynamics between morphological states (unicellular and multicellular) and reproductive traits (presence or absence of autospores and monospore/akinete formation), Bayesian analyses were conducted using BayesTraits v3.0.5 (Pagel et al. 2004) under a reversible-jump Markov chain Monte Carlo (MCMC) framework. Exponential priors with a mean of 10 were assigned to all transition rate parameters. To test whether the two traits evolved independently or dependently, marginal likelihoods were estimated using the stepping stone sampler under both the discrete independent and dependent models. The stepping stone analysis was performed using 100 stones, each run for 10,000 iterations. Log Bayes Factors were then calculated to evaluate support for the dependent model. MCMC chains were run for 1,000,000 generations with a burn-in of 10,000 generations. Convergence and mixing were assessed by ensuring effective sample sizes (ESS) > 200 using Tracer v1.7.2 (Rambaut et al. 2018). The analysis was performed on the same three-gene ML phylogram used for ancestral state reconstruction.

### Literature survey of Xanthophyceae taxonomy and character traits

To assess the current taxonomic status of traditionally recognized genera in Xanthophyceae, we used the treatment of Ettl (1978) as the primary reference framework. This was cross-checked against Silva (1979) to verify nomenclatural validity and correct potential inconsistencies. Current taxonomic status and availability of molecular sequence data were further evaluated using AlgaeBase (Guiry and Guiry 2026) and additional literature cited therein.

Morphological characters related to multicellularity and reproductive modes were compiled from Ettl (1978) and other primary taxonomic and life history studies for each genus or species. Following the framework of Knoll (2011), unicellular forms were defined as solitary or temporarily aggregated cells lacking persistent developmental integration, whereas multicellular forms were defined as filamentous or coenocytic bodies with stable physical connection, organism-level integration, or differentiated thallus organization. Zoospores are motile and typically uninucleate, whereas autospores, monospores, and akinetes are non-motile; synzoospores are motile but multinucleate propagules characteristic of siphonous taxa. Autospores are formed by internal cleavage into one or more daughter cells, monospores are single non-motile propagules formed without internal division, akinetes are also single-celled and transformed resting vegetative cells, and synzoospores contain multiple nuclei or coordinated cytoplasmic domains. Character codings and corresponding references are provided in Supplementary Table S2, while detailed criteria and explanatory notes are described in the Appendix.

## 3. Results

### Assembly and annotation

Transcriptome assemblies generated for 18 species yielded 12,989–37,597 clustered protein-coding ORFs across the sampled taxa, and their completeness assessed with BUSCO (eukaryota_odb10) ranged from the lowest in *Bumilleriopsis peterseniana* (55.3%) to the highest value observed in *Heterococcus viridis* (89.0%) (Supplementary Table S4). The majority of transcriptomes (12 out of 15) exhibited BUSCO completeness > 68.0%.

Among Xanthophyceae, chloroplast genomes ranged from 115,448 to 153,703 bp in size, with GC contents of 27.9–37.4%, and typically contained 137–153 CDS and 27–32 tRNA genes (Supplementary Table S5). Mitochondrial genomes ranged from 34,800 to 62,398 bp, with GC contents of 29.2–36.8%, and encoded 30–42 CDS and 23–42 tRNA genes (Supplementary Table S6). With the exception of *Pseudopleurochloris* sp. LCR-Ant16_1, whose chloroplast and mitochondrial genomes were each recovered as three contigs, all organelle genomes were assembled as complete circular molecules.

### Nuclear phylogeny and its validation

Both concatenation (260,437 amino acid sites, 680 partitions) and coalescent (680 gene trees) analyses of the nuclear dataset consistently resolved the ordinal-level diversification of Xanthophyceae in a stepwise, serial pattern, with maximal support across all nodes (100% UFBoot in the ML tree; local posterior probability = 1.0 in the coalescent tree; Fig. 2d,e). The topology recovered five successive divergences: (i) a basal clade comprising *Heterococcus* and *Chlorellidium*; (ii) a clade of *Pseudopleurochloris* strains; (iii) the Mischococcales clade (*Sphaerosorus* + *Botrydiopsis*); (iv) the Tribonematales clade (*Asterosiphon*, *Botrydium*, *Ophiocytium*, *Xanthonema*, *Bumilleriopsis*, *Tribonema*, *Pseudobumilleriopsis*, *Bumilleria*); and (v) Vaucheriales (*Vaucheria*).

In addition, familial relationships within Tribonematales, previously unresolved in a three-gene analysis (Maistro, Broady, et al. (2009); Fig. 2b), were robustly clarified. Within this order, *Asterosiphon* is recovered as sister to all remaining lineages, followed by *Botrydium* as sister to the remaining taxa, after which two well-supported sister clades are recovered: (*Ophiocytium*, (*Xanthonema*, *Bumilleriopsis*))) and (*Tribonema*, (*Pseudobumilleriopsis*, *Bumilleria*)), together forming a monophyletic group relative to *Botrydium*. Nearly all nodes within Tribonematales received maximal support in both analyses, with the exception of the node uniting these two derived clades, which showed slightly lower support (92% UFBoot) in the concatenation analysis.

All different ordinal-level topologies other than the main species tree were significantly rejected across all statistical tests of alternative topologies (i.e., KH, SH, AU, WKH, WSH tests), with *p*-values < 0.05 in every case (Table 1). This indicates that the alternative arrangements of these major lineages are strongly inconsistent with the signal in the concatenated dataset, and thus the inferred nested sister relationships among Xanthophyceae clades is robustly supported. Similarly, the traditional hypothesis grouping *Asterosiphon* and *Vaucheria* within a single order (e.g., Vaucheriales *sensu* Rieth (1980)), as well as the hypothesis uniting siphonous *Botrydium* with filamentous taxa such as *Tribonema*, *Pseudobumilleriopsis*, and *Bumilleria*, were also rejected (*p*-value < 0.05 in all tests). In contrast, the alternative topology constraining the monophyly of *Botrydium* + (*Ophiocytium*, *Xanthonema*, *Bumilleriopsis*) was not formally rejected, although it received only marginal support. In four out of five tests, the *p*-values were just above the rejection threshold (typically < 0.1 but > 0.05), indicating that this hypothesis is statistically weak and borderline incompatible with the data. This near-significant rejection suggests that, while not definitively excluded, the grouping is unlikely to reflect the true evolutionary relationship and is not strongly supported compared to the best tree.

**Table 1.**
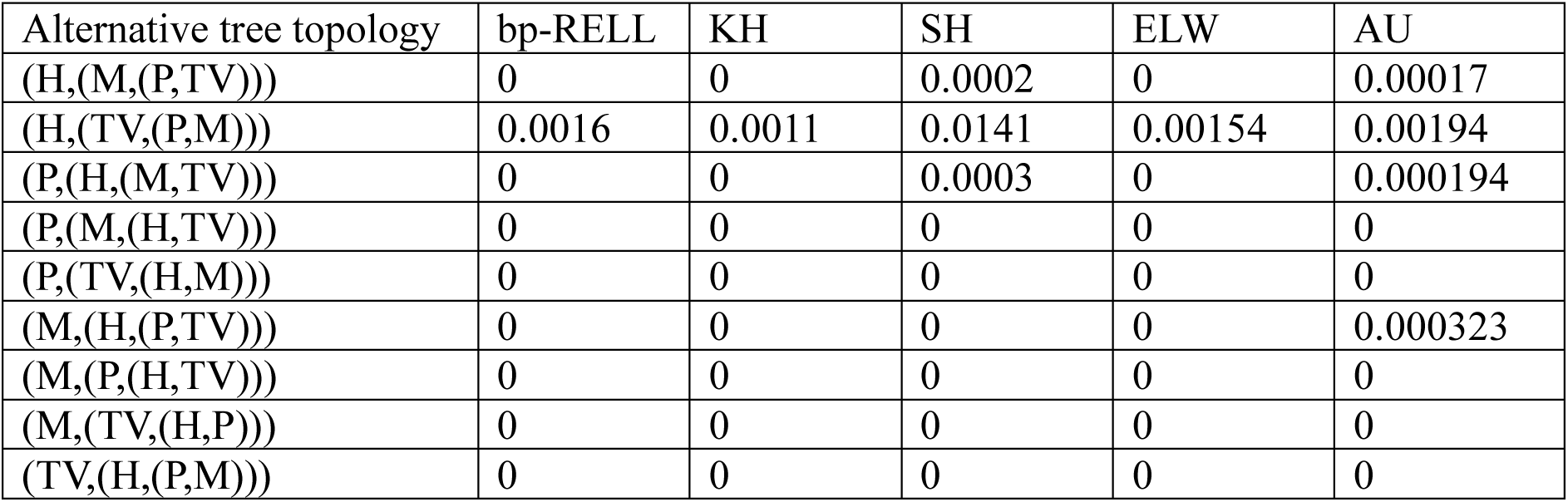
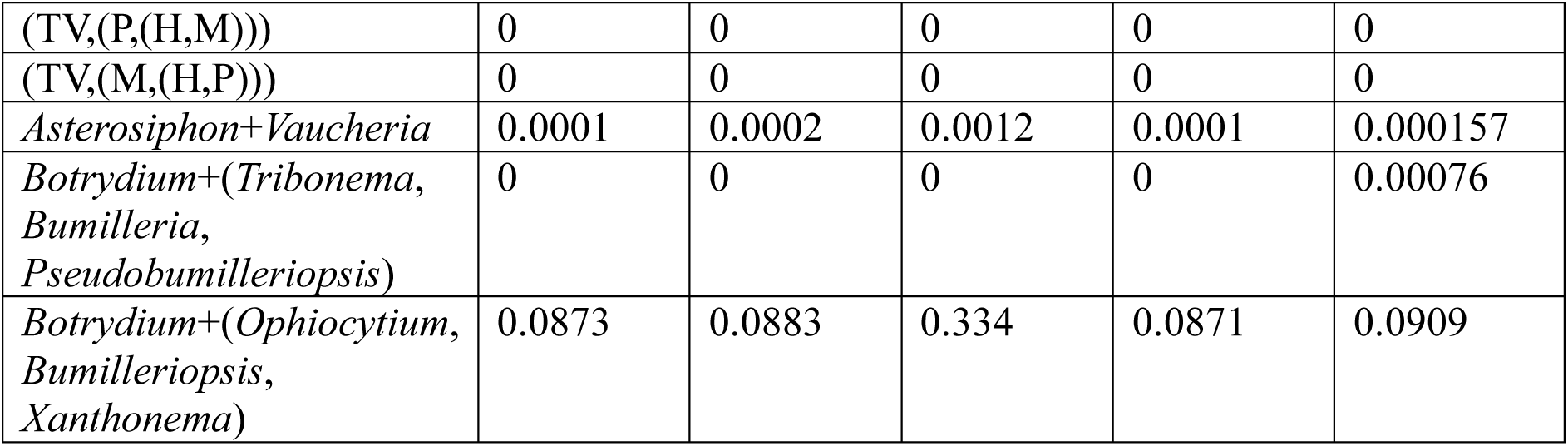
Statistics of alternative topology tests. All values shown are *p*-values. Values less than 0.0001 were treated as 0. bp-RELL, bootstrap proportion test using resampling estimated log-likelihoods; KH, weighted Kishino-Hasegawa test; SH, weighted Shimodaira-Hasegawa test; ELW, expected likelihood weight test; AU, approximately unbiased.; H, Heterococcales; P, Pseudopleurochloridales; M, Mischococcales; TV, Tribonematales+Vaucheriales.

| Alternative tree topology | bp-RELL | KH | SH | ELW | AU |
| --- | --- | --- | --- | --- | --- |
| (H,(M,(P,TV))) | 0 | 0 | 0.0002 | 0 | 0.00017 |
| (H,(TV,(P,M))) | 0.0016 | 0.0011 | 0.0141 | 0.00154 | 0.00194 |
| (P,(H,(M,TV))) | 0 | 0 | 0.0003 | 0 | 0.000194 |
| (P,(M,(H,TV))) | 0 | 0 | 0 | 0 | 0 |
| (P,(TV,(H,M))) | 0 | 0 | 0 | 0 | 0 |
| (M,(H,(P,TV))) | 0 | 0 | 0 | 0 | 0.000323 |
| (M,(P,(H,TV))) | 0 | 0 | 0 | 0 | 0 |
| (M,(TV,(H,P))) | 0 | 0 | 0 | 0 | 0 |
| (TV,(H,(P,M))) | 0 | 0 | 0 | 0 | 0 |
| (TV,(P,(H,M))) | 0 | 0 | 0 | 0 | 0 |
| (TV,(M,(H,P))) | 0 | 0 | 0 | 0 | 0 |
| *Asterosiphon*+*Vaucheria* | 0.0001 | 0.0002 | 0.0012 | 0.0001 | 0.000157 |
| *Botrydium*+(*Tribonema*,<br>*Bumilleria*,<br>*Pseudobumilleriopsis*) | 0 | 0 | 0 | 0 | 0.00076 |
| *Botrydium*+(*Ophiocytium*,<br>*Bumilleriopsis*,<br>*Xanthonema*) | 0.0873 | 0.0883 | 0.334 | 0.0871 | 0.0909 |

All gene tree metrics of nuclear genes, including branch support, root-to-tip CoV, stemminess, and tree length, are summarized in Supplementary Table S7. Phylogenies reconstructed from genes sorted by each metric are shown in Supplementary Fig. S1 (branch support), Fig. S2 (root-to-tip CoV), Fig. S3 (stemminess), and Fig. S4 (tree length). We tracked key ordinal (A-C in Fig. 3a) and familial (D-F in Fig. 3a) nodes to evaluate support changes across datasets.

**Figure 3.**
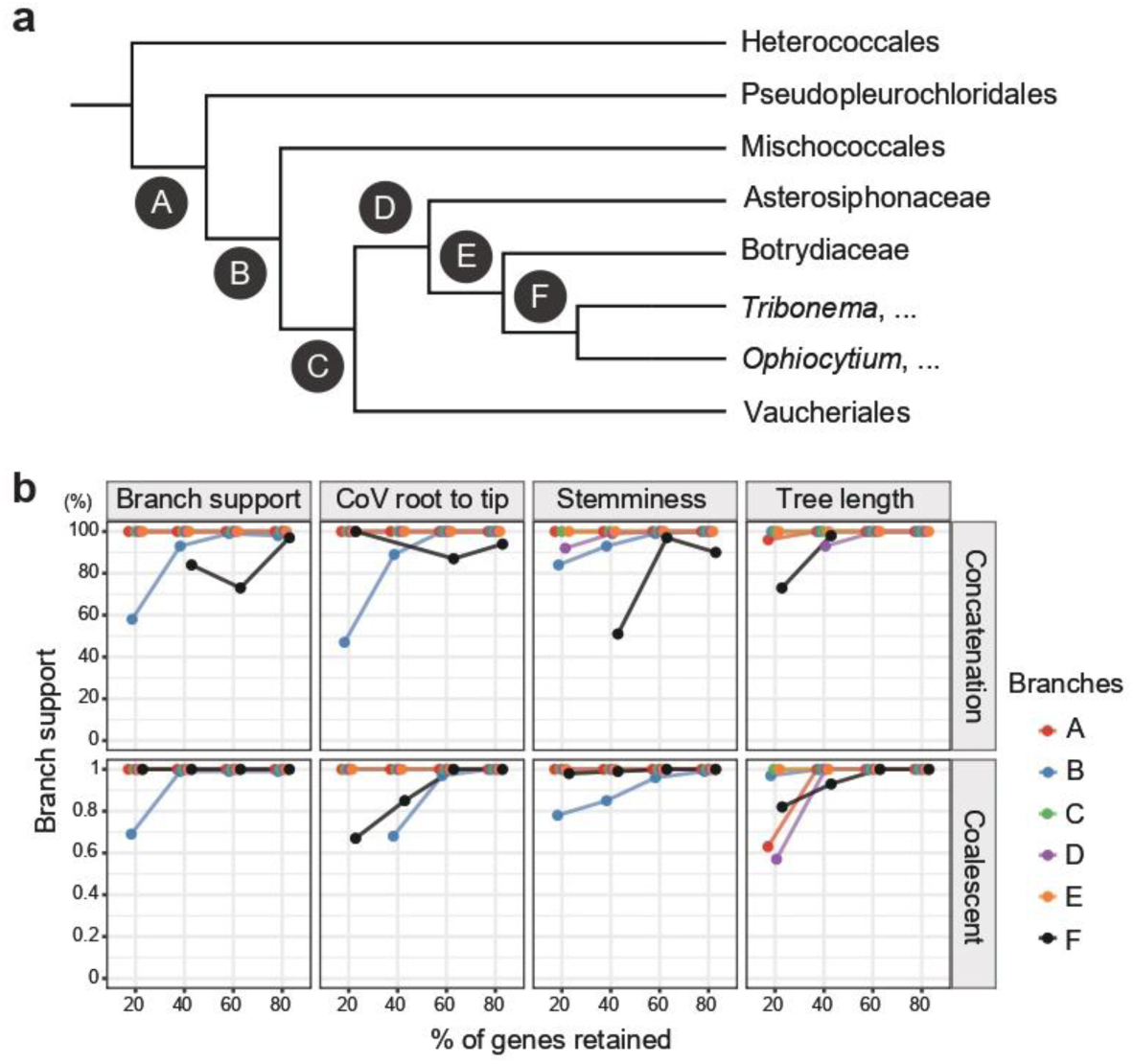
Species tree validation using gene-metric sorting. (a) Reference species tree based on nuclear phylogeny, with focal ordinal and familial nodes (A–F) indicated. (b) Changes in branch support at focal nodes as a function of the proportion of genes (%) retained under different sorting schemes. Points are omitted when alternative topologies inconsistent with the reference species tree are recovered.

Across all sorting schemes, branch support showed a consistent monotonic increase with the inclusion of higher proportions of genes (Fig. 3b). Node A, C, and E were nearly always maximally supported (100%) regardless of gene subset size or sorting metric. In contrast, node B (Mischococcales + (Tribonematales, Vaucheriales)) exhibited the progressive trend: for example, under root-to-tip CoV-based sorting, UFBoot support increased from 47% (20% genes) to 89% (40%), reaching 100% at ≥ 60% gene inclusion in concatenation analyses, with a similar pattern in coalescent trees (68% → 97% → 100%). A comparable trend was observed across other metrics, where node B support generally rose from ∼58–85% at 20% gene subsets to ∼98–100% at 80%.

Familial nodes (D-F) were overall more stable but still showed improvement with increasing gene sampling. Node D and E were consistently highly supported (mostly 100%), even at lower gene proportions. Node F (*Tribonema*,… + *Ophiocytium*,…) showed more variability at lower sampling (e.g., 67-82% in 20% coalescent datasets and 73% in tree length-sorted concatenation), but increased to ∼93-100% at ≥ 40-60% gene inclusion, reaching full support in most 60-80% datasets. Overall, these results demonstrate that increasing gene sampling consistently improves node support across both concatenation and coalescent frameworks, with particularly strong effects on moderately supported nodes (e.g., node B and F). Importantly, this monotonic trend is observed regardless of the gene-sorting metric, indicating that the inferred species phylogeny of Xanthophyceae is robust to gene selection strategy.

Correlation analysis of gene metrics (Supplementary Fig. S5) revealed that mean branch support was moderately negatively correlated with RF distance to the species tree (r = −0.55), indicating that loci with higher phylogenetic support tend to produce more congruent gene trees. Stemminess showed a weak positive correlation with RF distance (r = 0.19), whereas the coefficient of variation in root-to-tip distances (CoV) exhibited no meaningful relationship with RF distance (r ≈ 0.00). Tree length also showed only a weak association (r = −0.06). Among all variables, RF distance to the species tree was strongly correlated with mean RF distance to other gene trees (r = 0.91), suggesting that loci deviating from the species tree are consistently discordant across the dataset. Among explanatory variables, correlations were generally weak, although CoV showed a moderate negative correlation with tree length (r = −0.35), and mean branch support was moderately negatively correlated with mean RF distance to other gene trees (r = −0.55). Overall, these results indicate that mean branch support is the primary correlate of phylogenetic congruence, whereas CoV and tree length contribute little explanatory power, and stemminess captures a weak but independent aspect of gene-tree variation.

Multiple linear regression analysis revealed that gene-tree characteristics significantly explain variation in topological discordance to the species tree (F₄,₆₇₅ = 83.19, p < 2.2 × 10⁻¹⁶), with the model accounting for 33.0% of the variance (R² = 0.33). Among the predictors, mean branch support showed a strong negative association with RF distance (β = −0.0116, p < 2 × 10⁻¹⁶), indicating that loci with higher phylogenetic support tend to produce gene trees more congruent with the species tree. Stemminess exhibited a significant positive relationship with RF distance (β = 0.1623, p < 1 × 10⁻⁷), suggesting that tree shape also influences topological accuracy. In contrast, neither the coefficient of variation in root-to-tip distances (CoV; p = 0.215) nor total tree length (p = 0.108) showed significant effects.

### Organelle, taxon-rich three-gene phylogenies and taxonomic revisions

Chloroplast and mitochondrial phylogenies recovered fully congruent topologies at the class-and order-level (Fig. 4a,b). In the early divergences of Xanthophyceae and related lineages, both datasets strongly supported the monophyly of Chrysoparadoxophyceae, *Nematochrysis*, *Phaeobotrys*, and Xanthophyceae, each with maximal support, relative to Phaeothamniophyceae and the outgroup (Schizocladiophyceae + Phaeophyceae). Within this clade, Chrysoparadoxophyceae diverged first, followed by *Nematochrysis* (94% UFBoot in chloroplast; 100% in mitochondrial phylogeny), and then the *Phaeobotrys* + Xanthophyceae lineage (100% in both datasets).

**Figure 4.**
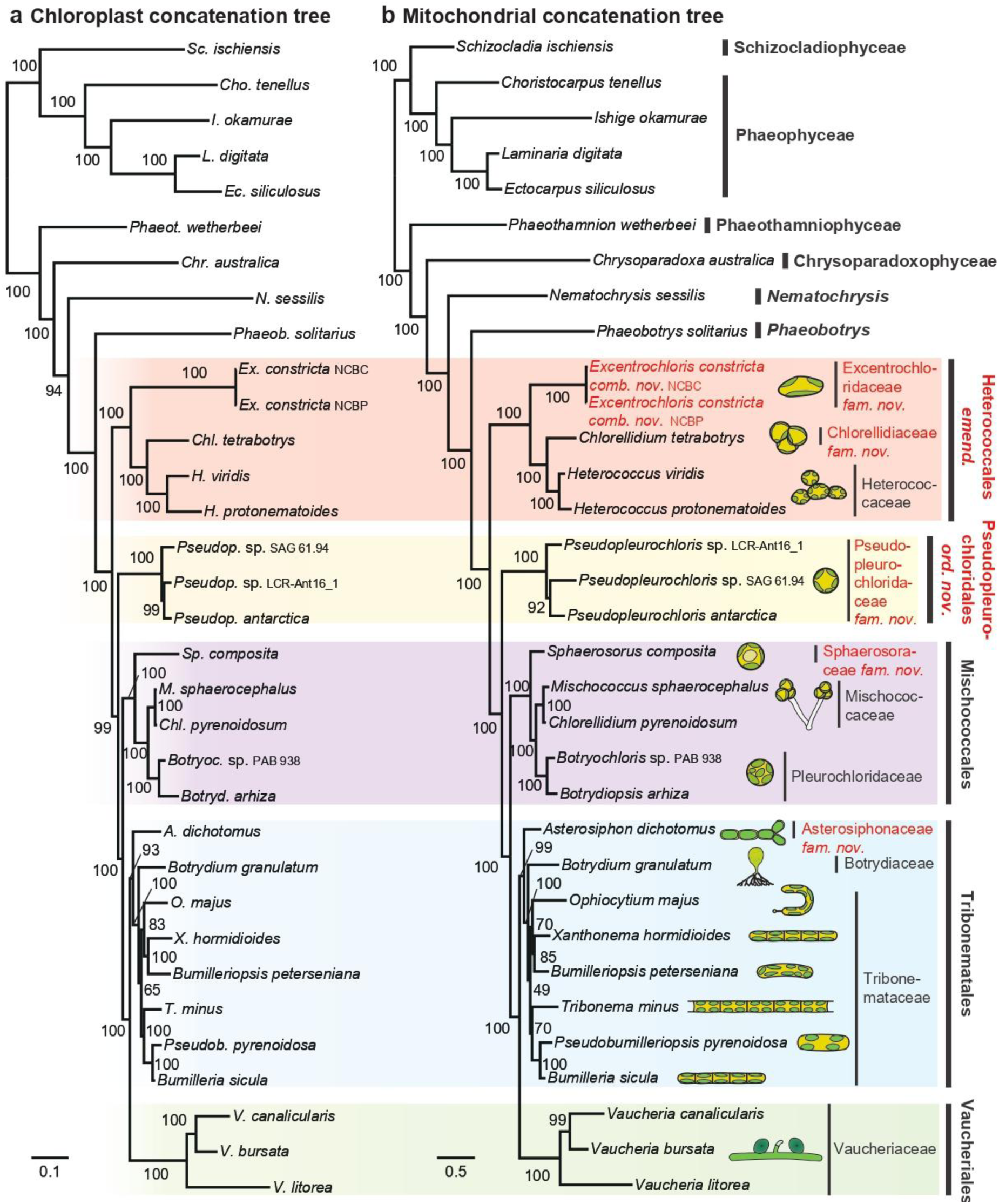
Organelle phylogenies of Xanthophyceae. (a) Chloroplast concatenation tree based on 141 genes and (b) mitochondrial concatenation tree based on 31 genes. Newly proposed taxonomic revisions are highlighted in red. Ultrafast bootstrap (UFBoot) values are shown on branches.

Order-level relationships were largely concordant with the nuclear phylogeny, while the organelle datasets provided expanded taxon sampling at the family level. The earliest-diverging lineage within Xanthophyceae comprised “*Botrydiopsis*” constricta (NCBC and NCBP strains), here treated as *Excentrochloris constricta* (Fig. 4), together with a fully supported clade of *Chlorellidium* and *Heterococcus*. The second lineage included three strains of *Pseudopleurochloris*, although their internal relationships differed between datasets: in the chloroplast tree, LCR-Ant16_1 grouped with *P. antarctica* (UFBoot = 99%), whereas in the mitochondrial tree, SAG 61.94 grouped with *P. antarctica* (UFBoot = 92%). The third divergence comprised *Sphaerosorus* and a clade including *Mischococcus* + “*Chlorellidium*” *pyrenoidosum* and *Botryochloris* + *Botrydiopsis*, all with maximal support in both organelle phylogenies. The fourth divergence included early-branching *Asterosiphon* and *Botrydium*, followed by two major clades: (i) *Ophiocytium* + (*Xanthonema*, *Bumilleriopsis*) and (ii) *Tribonema* + (*Pseudobumilleriopsis*, *Bumilleria*). Although this topology is consistent with the nuclear phylogeny, support for some nodes, particularly the *Ophiocytium* + *Tribonema* node, was reduced (<70% UFBoot) in both organelle datasets, likely reflecting limited phylogenetic signal. The final divergence corresponded to Vaucheriaceae, consistent with the nuclear tree.

The three-gene phylogeny (Supplementary Fig. S6) expanded species-level sampling within genera and generally supported their monophyly (e.g., *Excentrochloris*, *Chlorellidium* [except “*Chlorellidium*” *pyrenoidosum* PAB 785], *Heterococcus*, *Botrydiopsis*, *Xanthonema*, *Tribonema*, and *Vaucheria*). However, deeper (ordinal and familial) relationships were poorly resolved and often discordant with nuclear and organelle phylogenies, reflecting limited sequence information. Within the three-gene phylogeny, “*Botrydiopsis*” *constricta* (treated here as *Excentrochloris constricta*; Supplementary Fig. S6) strains NCBC and NCBP clustered closely with the type strain PAB 894 and the three of them formed a well-supported monophyletic group with *Excentrochloris fraunhoferiana*. This phylogenetic affinity is further supported by shared morphological characteristics among them, including (i) ellipsoidal cells, (ii) localized cell wall thickening, and (iii) cell constriction (Broady 1976; Begum 1999; Novis, Beer, et al. 2008; Hofbauer et al. 2011), all features consistent with those of the type species *Excentrochloris gigas* (Pascher 1939). Together, these lines of evidence provide a basis for the taxonomic revision proposed below.

*Excentrochloris constricta* (P. A. Broady) S.-W. Choi, P. A. Broady, P. M. Novis, R. A. Andersen et H. S. Yoon *comb. nov*.

Basionym: *Botrydiopsis constricta* P. A. Broady, *Br. Phycol. J*. 11: 388–392 (1976).

Comments: Distinguished from *Botrydiopsis* by its frequent transition to ellipsoidal to subcylindrical cell forms associated with constriction during vegetative division and localized cell wall thickening, whereas other species are predominantly spherical. Molecular phylogenetic analyses recover this species in a well-supported monophyletic clade with *Excentrochloris fraunhoferiana*, supporting its transfer to *Excentrochloris* and indicating concordance between morphological and phylogenetic evidence.

Order- and family-level taxonomy of Xanthophyceae can be established based on the robust phylogenetic framework inferred from nuclear, chloroplast, and mitochondrial datasets, as outlined below.

Class Xanthophyceae P. Allorge ex F. E. Fritsch 1935

Order 1 Heterococcales Pascher 1912 *emend.* S.-W. Choi, P. A. Broady, P. M. Novis, R. A. Andersen et H. S. Yoon

Cells or thallus unicellular, colonial, or filamentous; cells spherical, ellipsoidal, irregular or cylindrical; chloroplasts single to multiple, parietal; nuclei one to several per cell; reproduction by autospores, aplanospores, or zoospores; freshwater or terrestrial.

Comments: Pascher (1912) originally established Heterococcales as a morphologically defined group of coccoid, non-motile heterokont algae within his broader developmental classification of ‘Heterokontae’ (Heterokontophyta). Notably, this early circumscription did not center on the genus *Heterococcus*, but instead included a heterogeneous assemblage of coccoid forms defined by convergent morphology. Here, we emend Heterococcales to incorporate the phylogenetically defined lineage including the genus *Heterococcus*, thereby aligning the order with molecular systematics.

Family 1 Heterococcaceae P. C. Silva 1979

Genus Heterococcus

Family 2 Excentrochloridaceae S.-W. Choi, P. A. Broady, P. M. Novis, R. A. Andersen et H. S. Yoon *fam. nov*.

Cells initially spherical, ellipsoid, polyhedral, or pyriform; cell wall thin to thick, smooth, often thickened and stratified at one pole; chloroplasts numerous, discoid to polyhedral, sometimes reticulate; pyrenoids absent; cells solitary and initially uninucleate, becoming multinucleate; oil granules abundant; reproduction by autospores or zoospores; occurring in freshwater, moist soils among mosses and other vegetation, and on artificial substrates such as building walls.

Genus Excentrochloris

Family 3 Chlorellidiaceae S.-W. Choi, P. A. Broady, P. M. Novis, R. A. Andersen et H. S. Yoon *fam. nov*.

Cells coccoid, solitary or more commonly in pairs, tetrads, or irregular, often forming cushion-like, sessile masses on substrates; cells spherical to irregular, uninucleate or multinucleate; chloroplasts single to multiple per cell; reproduction by autospores or zoospores; freshwater or terrestrial.

Genus *Chlorellidium*

Order 2 Pseudopleurochloridales S.-W. Choi, P. A. Broady, P. M. Novis, R. A. Andersen et H. S. Yoon *ord. nov*.

Cells unicellular, coccoid, solitary or occasionally in small groups; chloroplasts parietal, multiple, lacking pyrenoid; plastid ultrastructure with thylakoids in triplets and a peripheral girdle lamella; nucleus single to multiple, associated with chloroplast endoplasmic reticulum; Golgi apparatus adjacent to nucleus; reproduction by autospores and zoospores; freshwater, terrestrial, or polar environments, including Antarctic regions.

Family Pseudopleurochloridaceae S.-W. Choi, P. A. Broady, P. M. Novis, R. A. Andersen et H. S. Yoon *fam. nov*.

Characters as for order Genus *Pseudopleurochloris*

Order 3 Mischococcales F. E. Fritsch 1927

Family 1 Sphaerosoraceae S.-W. Choi, P. A. Broady, P. M. Novis, R. A. Andersen et H. S. Yoon *fam. nov*.

Thallus colonial, forming regular spherical or ellipsoidal aggregates of few to many cells, often with tetrahedral or hollow organization; colonies without mucilage; cells spherical to ovoid, uniform in size, with smooth cell walls lacking ornamentation; chloroplasts parietal, discoid, one to several per cell, without pyrenoid; storage products absent as distinct oil bodies; reproduction by autospores or zoospores; resting stages present; freshwater, terrestrial, or marine.

Genus *Sphaerosorus*

Family 2 Mischococcaceae Pascher 1912

Genus *Mischococcus*

Family 3 Pleurochloridaceae Pascher 1937

Genus *Pleurochloris*, *Botryochloris*, *Botrydiopsis*

Comments: *Pleurochloris*, *Botryochloris* and *Botrydiopsis* were originally assigned to separate families, namely Pleurochloridaceae Pascher 1937, Botryochloridaceae Pascher 1937–1938 and Botrydiopsidaceae D. J. Hibberd 1980, respectively. However, our phylogenetic analyses recover *Pleurochloris*, *Botryochloris*, and *Botrydiopsis* as a strongly supported monophyletic group separated by very short internal branches, even relative to other familial-level divergences (Supplementary Fig. S6). Furthermore, the morphological characters historically used to distinguish these genera, such as the number of cells in colonial aggregates and individual cell size, are highly variable depending on environmental conditions and can vary substantially even within a single genus (e.g., *Botrydiopsis*; Ettl (1978)). These characters therefore do not appear to represent stable family-level distinctions. Accordingly, we treat these genera as belonging to a single family, Pleurochloridaceae, the oldest available family name for this clade.

Order 4 Tribonematales Pascher 1939

Family 1 Asterosiphonaceae S.-W. Choi, P. A. Broady, P. M. Novis, R. A. Andersen et H. S. Yoon *fam. nov*.

Thallus siphonous and coenocytic, dichotomously branched, forming radial rosette-like systems on substrate; reproduction by internal cyst formation within siphons, cysts germinating into new thalli or producing amoeboid or walled spores; freshwater or terrestrial.

Genus *Asterosiphon*

Family 2 Botrydiaceae Rabenhorst 1863

Genus *Botrydium*

Family 3 Tribonemataceae G. S. West 1904

Genus Tribonema, Xanthonema, Ophiocytium, Heterothrix, Bumilleria, Bumilleriopsis, Pseudobumilleriopsis

Order 5 Vaucheriales Blackman et Tansley 1902

Family Vaucheriaceae Dumortier 1822

Genus Vaucheria

### Ancestral character states and transition dynamics

All multicellularity and reproductive-mode character states included in the character evolution analyses, along with the literature sources used for character coding, are listed in Supplementary Table S2.

Ancestral character states were inferred using phytools under an equal-rates (ER) model based on the best Akaike Information Criterion (AIC) score and are shown in Fig. 5a. The results obtained under the all-rates-different (ARD) model are also presented in Supplementary Fig. S7, showing that the major ancestral state predictions at each node were consistent with those inferred under the ER model, differing only in the posterior probability proportions. Under the ER model, deep ancestral nodes within Xanthophyceae and related lineages were predominantly inferred as unicellular. These include the common ancestor of Xanthophyceae and Phaeothamniophyceae (posterior probability: unicellular 0.829, multicellular 0.171), the common ancestor of Xanthophyceae (unicellular 0.995, multicellular 0.005), the ancestor of Pseudopleurochloridales + Mischococcales (unicellular 0.994, multicellular 0.006), and the ancestor of Mischococcales + Tribonematales (unicellular 0.983, multicellular 0.017). In contrast, the Tribonematales + Vaucheriales node showed a predominantly multicellular ancestral state prediction (unicellular 0.184, multicellular 0.816).

**Figure 5.**
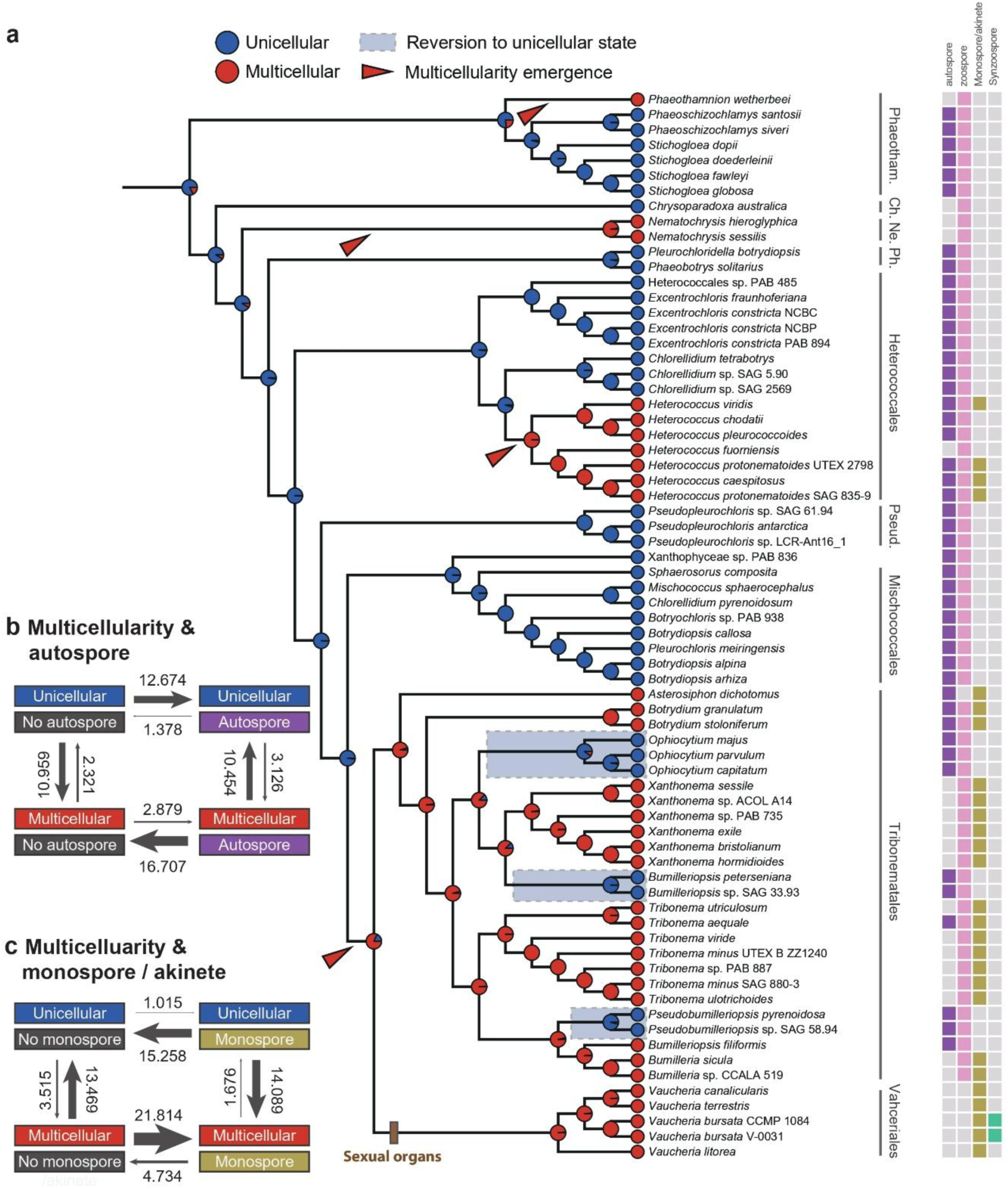
Morphological and reproductive evolution in Xanthophyceae. (a) Ancestral state reconstruction of multicellularity inferred using phytools. Posterior probabilities for each character state are shown at internal nodes (unicellular, blue; multicellular, red). The backbone phylogeny is presented as a cladogram. Phaeotham., Phaeothamniophyceae; Ch., Chrysoparadoxophyceae; Ne., *Nematochrysis*; Ph., *Phaeobotrys*; Pseud., Pseudopleurochloridales. Presence or absence of reproductive characters in each species is indicated on the right side of the tree (autospore, purple; zoospore, pink; monospore/akinete, dark yellow; synzoospore, cyan). Bayesian estimates of transition rates inferred using BayesTraits are shown for correlated evolution between multicellularity and the presence/absence of (b) autospores and (c) monospores or akinetes. Arrows indicate the direction of transitions, with arrow widths proportional to the estimated transition rates.

When considering the ancestral nodes of individual lineages, Phaeothamniophyceae (unicellular 0.767), *Phaeobotrys* + *Pleurochloridella* (unicellular 0.976), Heterococcales (unicellular 0.986), Pseudopleurochloridales (unicellular 1.000), and Mischococcales (unicellular 0.998) were inferred to have predominantly unicellular ancestors. In contrast, *Nematochrysis* (multicellular 0.978), Heterococcaceae (multicellular 0.998), Tribonematales (multicellular 0.949), and Vaucheriales (multicellular 0.983) were inferred to have predominantly multicellular ancestors. Four independent origins of multicellularity were inferred in *Phaeothamnion*, *Nematochrysis*, *Heterococcus*, and the common ancestor of Tribonematales + Vaucheriales (red arrowheads in Fig. 5a). In contrast, reversions from multicellular to unicellular states were inferred at three nodes within Tribonematales: from the *Xanthonema* + *Ophiocytium* node (multicellular 0.876) to *Ophiocytium* (unicellular 0.906), from the *Xanthonema* + *Bumilleriopsis* node (multicellular 0.877) to *Bumilleriopsis* (unicellular 0.991), and from the *Pseudobumilleriopsis* + *Bumilleria* node (multicellular 0.965) to *Pseudobumilleriopsis* (unicellular 0.959; blue boxes in Fig. 5a).

For both trait combinations analyzed in BayesTraits, the dependent model yielded substantially higher marginal likelihoods than the independent model, with Log Bayes Factor values indicating strong to very strong support for correlated character evolution (autospore: 9.994; monospore/akinete: 12.4564), following the criteria of Kass and Raftery (1995).

For analyses incorporating autospore presence or absence, transition rates differed markedly depending on the morphological state (Fig. 5b). Under the unicellular condition, the transition toward autospore presence (12.674) was more than nine times higher than the reverse transition (1.378). In contrast, under the multicellular condition, the transition toward autospore absence (16.707) was more than five times higher than the reverse transition (2.879). When the reproductive state was fixed as autospore absence, the transition from unicellularity to multicellularity (10.959) was more than four times higher than the reverse transition (2.231). Conversely, when the autospore cycle was present, the transition from multicellularity to unicellularity (10.454) was approximately three times higher than the reverse transition (3.126)

In a similar manner, analyses incorporating the presence or absence of monospores/akinetes also showed transition dynamics that varied according to multicellular state, but in the opposite direction compared with the autospore pattern (Fig. 5c). Under the unicellular condition, transitions toward monospore/akinete absence (15.258) were approximately 15 times more frequent than transitions toward presence (1.015). In contrast, under the multicellular condition, transitions toward monospore/akinete presence (21.814) were more than four times more frequent than transitions toward absence (4.734). When the reproductive state was fixed, the inferred transition dynamics also differed markedly. Under the monospore/akinete absence condition, transitions from multicellularity to unicellularity (13.469) were more than three times more frequent than transitions from unicellularity to multicellularity (3.515). Conversely, under the monospore/akinete presence condition, transitions from unicellularity to multicellularity (14.089) were approximately eight times more frequent than the reverse transition from multicellularity to unicellularity (1.676).

### Literature survey of taxonomic status

All recent cases of taxonomic revision since Ettl (1978) are summarized in Supplementary Table S8 and Supplementary Fig. S8. Among the 115 genera recognized by Ettl, 87 (75.7%) lack any sequence data. Eleven genera (9.6%) have been transferred from Xanthophyceae to other lineages, most frequently to Eustigmatophyceae (6 genera, 5.2%). Only 17 genera (14.8%) have been confirmed with sequence data, either reassigned to a different family within Xanthophyceae (7, 6.1%) or retained in their original family (10, 8.7%).

Across orders of Xanthophyceae, no genera in Chloramoebales (flagellate), Rhizochloridales (amoeboid), or Heterogloeales (palmelloid) have been confirmed as Xanthophyceae based on sequence data (Supplementary Fig. S8). Instead, Rhizochloridales includes transfer cases to Synchromophyceae and Chlorarachniophyta, while Heterogloeales includes *Pleurochloridella* (incertae sedis, sister to Xanthophyceae) and transfers to Eustigmatophyceae. In Mischococcales, five genera have been transferred to Eustigmatophyceae, along with single transfers to Trebouxiophyceae and Oomycota. In contrast, no transfer events to non-xanthophycean lineages were detected in Tribonematales, Botrydiales, or Vaucheriales, all of which comprise multicellular representatives.

## 5. Discussion

### Emergence of multicellularity and reproductive changes in Xanthophyceae

The relationship between multicellularity and reproductive strategy has been extensively explored in theoretical models, yet empirical studies remain limited and strongly biased toward a small number of model systems (Ratcliff, Herron, et al. 2013; Brunet and King 2017). The present analyses show that Xanthophyceae provides an empirical system in which repeated changes in body organization can be examined together with shifts in asexual reproductive strategy. Because the genome-scale phylogeny established here resolves major xanthophycean relationships with strong support, these transitions can be interpreted in a robust comparative framework rather than as isolated morphological observations.

A central result of this study is that the emergence of multicellularity was not necessarily associated with unicellular (uninucleate) propagules, as proposed by the “unicellular bottleneck” theory (Grosberg and Strathmann 1998). Instead, ancestral character estimation and Bayesian transition analyses showed that multicellular lineages were more strongly associated with the loss of autospore-based reproduction and the gain of monospore-or akinete-type propagules (Fig. 5). This pattern is notable because autospores consist of multiple small uninucleate daughter cells produced through internal cleavage, whereas monospores and akinetes generally represent single, larger, non-motile propagules that may retain a multinucleate condition. These results suggest that multinucleate propagules may confer ecological or evolutionary advantages despite the potential disadvantages associated with increased genetic heterogeneity.

The ecological and evolutionary importance of multinucleate propagules may lie in their role as large, resource-rich units that enhance establishment under variable environmental conditions. In Xanthophyceae, most lineages reproduce predominantly asexually, with only a few genera, such as *Vaucheria* and *Botrydium*, frequently expressing sexual organs (Ettl 1978; Rieth 1980; Maistro, Broady, et al. 2017). In this context, monospore- and akinete-type propagules may have provided particular advantages in freshwater and terrestrial habitats, where post-dispersal survival is likely to be more environmentally unstable and condition-dependent than in many marine systems (Steele et al. 2019). By providing greater size, stored resources, and resistance to unfavorable conditions, these larger and often multinucleate propagules may have increased the success of asexual reproduction in multicellular lineages (Simberloff 2009). The xanthophycean pattern is therefore consistent with the view that unicellular propagation is not an obligatory prerequisite for the evolution of multicellularity, but rather one of several reproductive strategies that may later become advantageous in lineages where tighter control of genetic heterogeneity is favored (Ratcliff, Herron, et al. 2013; Pichugin et al. 2017).

Although multinucleate propagules may retain lingering effects of genetic heterogeneity within the organism, xanthophytes may possess several intrinsic mechanisms that mitigate such heterogeneity. First, nuclear selection (intraorganismal selection among nuclei within a shared cytoplasm) may reduce the contribution of deleterious nuclear variants prior to propagule formation (Otto and Orive 1995). Such nuclear-level filtering has been discussed in relation to multinucleate fungal heterokaryons and variegated plants (Stewart 1978; Buss 1983; Samils et al. 2014) and may be particularly relevant in coenocytic or siphonous algae, where numerous nuclei coexist within an integrated cytoplasmic system before being partitioned into reproductive structures.

Second, multinucleate organization itself may buffer the phenotypic effects of deleterious mutations. In a shared cytoplasm, defective nuclei may be masked by functional nuclei (Otto and Whitton 2000), allowing large propagules to develop successfully even without a strict single-nucleus bottleneck. This may be particularly relevant in coenocytic taxa such as *Vaucheria*, *Botrydium*, and *Asterosiphon*, where organismal integration is achieved without regular cellular compartmentalization. In such lineages, the immediate advantage of a large, resource-rich propagule may outweigh the cost of transmitting multiple nuclei.

Third, the evolution of localized cleavage, septation, or reproductive compartmentalization may have provided an intermediate solution between unrestricted multinuclearity and a strict unicellular bottleneck. Even when vegetative cells are multinucleate, filamentous tribonematalean species undergo cell division before reproduction, and in some cases additional divisions during the formation of monospores or akinetes, thereby reducing the number of nuclei transmitted to each propagule (Lokhorst and Star 1988; Massalski et al. 2009). This would not eliminate genetic heterogeneity entirely, but it could limit the nuclear pool contributing to each offspring while preserving the developmental advantages of larger propagules. Such a ‘partial bottleneck’ may have been sufficient for simple multicellular xanthophytes, especially in lineages where asexual reproduction remained the dominant life history strategy.

Finally, sexual reproduction may serve as an additional backup mechanism for controlling nuclear heterogeneity in some multicellular lineages. *Vaucheria* is a notable example because it combines large asexual propagules, including monospores (aplanospores) and multinucleate synzoospores, with a specialized oogamous sexual cycle (Ott and Brown Jr 1974). During sexual reproduction, many nuclei can enter developing reproductive structures, but only one nucleus is ultimately retained in the oogonium and spermatozoid (Gross 1937; Ott and Brown Jr 1978). This suggests that *Vaucheria* may use sexual development as a stronger bottleneck while retaining large multinucleate propagules for asexual reproduction. How nuclei are selectively retained or eliminated during this process remains unclear and should be a key target for future genomic and developmental studies.

Overall, the evolution of multicellularity in Xanthophyceae involved not only changes in body organization, but also repeated restructuring of asexual reproductive strategies. The association between multicellularity and monospore- or akinete-type propagules suggests that large, often multinucleate propagules were not evolutionary anomalies, but recurring components of xanthophycean multicellular life histories. These findings broaden the discussion of multicellular evolution by showing that continuity of stable multicellular lineages does not necessarily depend on a strict unicellular bottleneck; rather, different propagule strategies may be favored according to environmental context, reproductive mode, and lineage-specific developmental architecture.

### Phylogeny-based higher-level taxonomy of Xanthophyceae

The phylogenomic framework presented here provides a decisive improvement in resolving higher-level relationships within Xanthophyceae, particularly at the ordinal and familial levels. Previous multigene studies were able to identify major clades but lacked sufficient phylogenetic signal (three genes or less) to robustly resolve deep divergences, resulting in weak support and inconsistent topologies across datasets (Maistro, Broady, et al. 2009; Rybalka et al. 2020). In contrast, the genome-scale nuclear dataset (680 genes), together with plastid and mitochondrial data, yields a fully resolved backbone phylogeny, with complete concordance between concatenation and coalescent approaches and no detectable conflict among genomic compartments. This level of resolution allows, for the first time, the clear identification of five successive ordinal divergences and the robust delimitation of familial relationships, particularly within historically problematic groups such as Tribonematales (Maistro, Broady, et al. 2017).

However, a large proportion of genera historically assigned to Xanthophyceae have remained uncharacterized at the molecular level, with many lacking sequence, pigment, or ultrastructural data since their original descriptions (Pascher 1939; Ettl 1978). While this incomplete sampling is substantial, it does not preclude higher-level taxonomic revision. Rather, it underscores the limitations of traditional morphology-based classifications, which are now recognized as highly homoplastic across heterokontophytes and other protist lineages (Andersen 2004; Cavalier-Smith and Chao 2006). Characters such as unicellular flagellate organization, palmelloid or coccoid growth, and amoeboid morphology recur across distantly related groups, rendering them unreliable indicators of phylogenetic affinity (Yang, Boo, et al. 2012). Consistent with this, our ancestral character reconstruction indicates repeated re-emergence of Mischococcales-like unicellular coccoid forms within Xanthophyceae (Fig. 5), suggesting that such morphologies are evolutionarily labile and prone to convergence.

This homoplasy is further illustrated by the historical reassignment of morphologically “xanthophycean” taxa to other lineages, including Eustigmatophyceae and additional heterokont groups. Notably, Eustigmatophyceae, originally segregated from Xanthophyceae based on cytological and ultrastructural evidence (Hibberd and Leedale 1970; Hibberd 1981), has received the largest number of transferred genera among related lineages even after the initial separation of the class (six genera; Amaral et al. (2021); Barcytė et al. (2022); Supplementary Table S8 and Fig. S8). These taxa often share ecological and morphological similarities with xanthophytes, including terrestrial or freshwater habitats, green pigmentation with reduced or absent chlorophyll c, the presence of accessory pigments such as violaxanthin or vaucheriaxanthin, and simple coccoid morphologies. However, subsequent ultrastructural studies demonstrated that these similarities are misleading, and that eustigmatophytes possess distinct diagnostic features, including a reduced flagellar apparatus and an extraplastidial eyespot, supporting their recognition as a separate class (Hibberd and Leedale 1970; Hibberd and Leedale 1972). This example highlights the extent to which traditional circumscriptions of Xanthophyceae might have been shaped by convergent ecological and biochemical traits rather than shared ancestry.

In contrast, phylogenetic approaches have consistently provided more stable and reproducible frameworks for higher-level classification across eukaryotes (Burki et al. 2020). Within Heterokontophyta specifically, sequence-based phylogenetic analyses have repeatedly led to major revisions and the recognition of novel class- and order-level lineages, often overturning assumptions derived from homoplastic, morphology-based taxonomy. These include the coccoid and filamentous Phaeothamniophyceae, formerly classified within Chrysophyceae (Bailey et al. 1998; Graf et al. 2020a); the flagellate Olisthodiscophyceae, previously placed in Raphidophyceae (Barcytė et al. 2021); the colonial and parenchymatous Phaeosacciophyceae, distinct from parenchymatous brown algae (Graf et al. 2020b); and the amoeboid Synchromophyceae (Grant et al. 2009; Schmidt et al. 2015), including taxa such as *Chlamydoxa* previously assigned to Xanthophyceae (Horn et al. 2007). These revisions, together with broader refinements to heterokont phylogeny (Yang, Boo, et al. 2012; Derelle et al. 2016; Terpis et al. 2025), demonstrate the limitations of morphology-based classification at higher taxonomic ranks. In this context, the absence of molecular data for many described genera should not be taken as justification for retaining historically broad and potentially artificial taxa. Instead, we propose a conservative, phylogeny-based revision that recognizes only those orders and families robustly supported by genomic-scale data. Our classification is therefore best interpreted as a backbone framework for sequenced diversity, rather than a definitive placement of all described genera, with unsampled taxa treated as provisionally assigned pending molecular characterization.

It is notable that no comprehensive order- or family-level revision of Xanthophyceae has been undertaken since early multigene studies, which yielded only loosely defined informal designations (e.g., Botrydiopsalean, Chlorellidialean, Tribonematalean, and Vaucherialean clades *sensu* Maistro, Broady, et al. (2009)). The phylogenomic framework presented here, supported by extensive gene sampling and strong concordance across various genomic data and analytical approaches, provides a timely and necessary update. By prioritizing well-supported clades in accordance with ‘phylogenetic systematics’ (De Queiroz and Gauthier 1994; Hennig 1999), while explicitly acknowledging remaining uncertainty, this approach establishes a stable and testable foundation for future systematic and evolutionary studies of Xanthophyceae.

## Acknowledgements

We thank Wolf-Henning Kusber (Freie Universität Berlin) for valuable discussions on taxonomic issues, Jihoon Jo (Honam National Institute of Biological Resources) for insightful discussions on Xanthophyceae, Hocheol Kim (Sungkyunkwan University) for assistance with *Vaucheria* sampling, and Jiwoo Lee and Sihoon Lee (Gyeonggi Science High School) for their assistance with organelle genome assembly.

## Data availability

The sequence data and genomes have been deposited in NCBI under BioProject PRJNA1506367. Accession numbers for all newly sequenced 18S rRNA sequences and organelle genomes are provided in Supplementary Table S2, S5 and S6. The sequence alignments, phylogenetic trees, and genome datasets used for the phylogenomic analyses in this study are available from Dryad (DOI: 10.5061/dryad.9w0vt4bxf).

## Appendix. Definitions of multicellularity and propagule characters in Xanthophyceae and related heterokontophytes

This appendix of definitions is not intended to replace traditional terminology used for heterokontophytes, but rather to provide clearer meanings for evolutionary analyses of character transitions in this study. It aims to minimize ambiguity arising from inconsistent or ambiguous definitions and thereby reduce unnecessary confusion in comparative framework. Moreover, character states were documented based on available observations of their presence or absence in each genus or species. Therefore, absence of evidence should be interpreted with caution and should not be taken as definitive evidence that a character is biologically absent, particularly when the phenotype may be rarely expressed, condition-dependent, or insufficiently documented.

### 1. Conceptual framework for multicellularity

Multicellularity is treated here as a condition of persistent biological integration rather than merely the presence of more than one nucleus or more than one cell in proximity. In the sense discussed by Knoll (2011), simple multicellularity includes filaments, sheets, clusters, or other reproducible bodies that arise through cell division and retain physical connection among daughter cells, sometimes with basic somatic and reproductive differentiation. Complex multicellularity involves stronger functional integration, including intercompartmental communication, developmentally regulated differentiation, and mechanisms that overcome diffusion limits in larger bodies. In both cases, a species is regarded as multicellular only when it exhibits a tightly maintained physical association among cells or compartments together with some degree of cytological integration.

In Xanthophyceae, multinucleate organization is widespread and may represent a common cytological condition rather than multicellularity by itself (Maistro, Broady, et al. 2017). Many xanthophycean cells begin as uninucleate juvenile cells and become multinucleate before cell division, zoospore formation, autospore formation, or other reproductive events. Therefore, multinuclearity alone is not sufficient evidence for multicellularity in this group. A multinucleate solitary cell such as *Botrydiopsis*, *Ophiocytium*, or *Excentrochloris*, even though they form a kind of ‘elongated’ form in a single cell level, should not be treated as multicellular unless additional evidence shows an integrated multicellular body plan, persistent regional differentiation, or coenocytic thallus-level organization, following criteria suggested by Knoll (2011).

Unicellular organization is defined here as a solitary, coccoid, ellipsoid, rod-like, or otherwise single-cell condition in which cells may be free-living, mucilage-embedded, temporarily grouped, or retained within the parental wall after reproduction, but do not form an obligate serial body with persistent developmental integration. Colonial or clumped forms are therefore not regarded as multicellular in the present framework.

Multicellular organization is a filamentous or coenocytic developmental condition in which cellular components are integrated into a stable body plan rather than existing as solitary or accidental aggregates. Filamentous multicellularity is defined by growth in which daughter cells remain physically connected as repeated serial units, producing persistent chains, branched systems, or reproducible pseudofilamentous thalli, as seen in *Tribonema*-type and *Heterococcus*-type taxa. The critical feature is not the number of nuclei within any one cell, but the necessary physical connection of multiple cellular units within a stable and repeated developmental architecture. Coenocytic or siphonous taxa are likewise regarded as multicellular-grade organisms when they form integrated macroscopic or differentiated thalli despite lacking regularly repeated cross walls. In genera such as *Vaucheria*, *Botrydium*, and *Asterosiphon*, multicellularity is supported by (i) organ or regional differentiation (e.g., rhizoids, akinetes, and sexual organs), (ii) long-distance cytoplasmic and nuclear transport, (iii) localized wall cleavage or septation during reproduction, injury response, or organ formation, and (iv) large macroscopic size, all of which indicate an integrated organismal body plan rather than a simple multinucleate unicell.

### 2. Propagule terminology

The terminology for propagules follows Pascher (1925) as far as possible, but with explicit caveats because many older xanthophycean descriptions used the terms aplanospore, autospore, resting spore, and akinete in overlapping or inconsistent ways. The distinctions adopted here are therefore based primarily on developmental origin rather than on morphology alone, with additional consideration given to (i) nuclear condition (i.e., uninucleate or multinucleate) and (ii) the occurrence or absence of internal cell divisions within the mother cell prior to dispersal.

A zoospore is a motile uninucleate propagule bearing flagella. In xanthophycean and related heterokont lineages, zoospores typically possess two unequal flagella, although the shorter flagellum was often overlooked or difficult to observe in older literature. Zoospores may be produced singly, in pairs, or in larger numbers depending on the genus and species.

An autospore is a non-motile uninucleate daughter cell produced after the protoplast undergoes internal cleavage within the mother cell wall, usually yielding one or more reduced daughter units that are immediately capable of vegetative growth. The essential criterion is subdivision of the original protoplast into separate daughter cells. Accordingly, when several small non-motile bodies are formed within a single mother cell through repeated internal divisions, even if described historically as “aplanospores” or by other names, they are interpreted here as autospores.

A monospore is used here for a single non-motile endogenous propagule formed without a flagellated stage and without further internal division of the protoplast inside the mother cell wall. Although the term has been widely used in red algae and fungi, it has not been applied in xanthophytes. In Pascherian usage, many instances of a single internally formed “aplanospore” in filamentous xanthophytes correspond closely to this concept. Such propagules are often more resistant than autospores, may possess a thicker wall, and may germinate only after a resting interval. The practical distinction between autospore and monospore used here is therefore the presence or absence of internal cleavage into multiple daughter units, while biological interpretation should also consider wall formation and dormancy behavior.

An akinete is a resting cell formed by direct transformation of an existing vegetative cell. Unlike autospores or monospores, which arise endogenously within a mother cell, an akinete is essentially the original vegetative cell converted into a thick-walled resistant stage that resumes growth or reproduction after dormancy. In filamentous taxa, akinetes may remain in chains or occur as filament akinetes. For analytical purposes, monospores and akinetes may overlap functionally because both usually represent a single propagule body formed without multiple internal divisions, often approximating the size of the original cell, frequently entering germination already in a multinucleate state, and commonly showing correlated occurrence in multicellular xanthophytes (i.e., both present or both absent). For this reason, they are treated as a single character state in Bayesian analyses of propagule evolution.

A synzoospore is a compound motile propagule, especially characteristic of siphonous forms such as *Vaucheria*. It contains multiple nuclei or coordinated cytoplasmic domains and reflects the integrated multinucleate organization of the parent thallus. It should therefore be distinguished from ordinary uninucleate zoospores.

### 3. Genus-level summaries

The following summaries emphasize only body organization, multicellularity-relevant traits, and propagules. They are intentionally concise and should be read as descriptive statements rather than strict numerical scores. These accounts are based primarily on Ettl (1978), Rieth (1980), Hibberd (1990), Maistro, Broady, et al. (2017), Guiry and Guiry (2026), among others; more detailed generic descriptions can be found in those references.

#### Phaeothamnion

*Phaeothamnion* is a filamentous genus with branched vegetative filaments and is therefore treated as a multicellular filamentous organization. Vegetative filaments may transform into palmelloid stages in older cultures. Zoospores are produced, and their occurrence supports a life cycle that alternates between filamentous growth and motile dispersal.

#### Phaeoschizochlamys

*Phaeoschizochlamys* is a palmelloid or mucilage-associated form in which cells occur singly or in small groups of two to four. It is best described as colonial rather than true multicellular because the grouped state lacks the persistent serial developmental integration of a filament. Cells form two or four autospores, and zoospore can also be produced.

#### Stichogloea

*Stichogloea* forms gelatinous colonies in which cells commonly occur as tetrads or short chains within mucilage. This organization is colonial and should not be equated with filamentous multicellularity unless stronger developmental integration is demonstrated. Reproduction includes autospores, and flagellate zoospore cells have been reported.

#### Chrysoparadoxa

*Chrysoparadoxa* is a small unicellular benthic heterokont with a walled attached vegetative stage. Each vegetative cell gives rise to a single naked zoospore, and daughter cells may briefly become amoeboid before final attachment. It is therefore treated as unicellular.

#### Nematochrysis

*Nematochrysis* is a filamentous marine heterokont and is treated as multicellular in the broad filamentous sense when it forms persistent threads. Motile reproductive stages were documented.

#### Phaeobotrys

*Phaeobotrys* is a solitary coccoid. Small young cells grow into larger thick-walled multinucleate cells with numerous chloroplasts, but this multinuclear enlargement is not multicellularity. Reproduction includes 8 to 32 autospores and *Ochromonas*-like zoospores.

#### Pleurochloridella

*Pleurochloridella* is a free-living unicellular genus in which cells may remain temporarily grouped within the maternal wall after division. Such retention is not considered true multicellularity because the cells do not form a persistent integrated filament or thallus. Zoospores and autospores are reported, and autospores may be formed in pairs or rarely in fours.

#### Heterococcus

*Heterococcus* is best treated as a simple multicellular or pseudofilamentous lineage when the organism forms branched or compact systems of physically connected cells. Zoospores, autospores, and akinetes or akinete-like resting cells are reported in several species, including taxa with terrestrial or Antarctic habitats.

#### Chlorellidium

*Chlorellidium* forms tetrads, small groups, and larger cushion-like aggregates that are often held together by remnants of maternal cell walls. These aggregates are best described as colonial or pseudocolonial rather than fully multicellular because integration is mainly through retained walls and close packing. Reproduction is by autospores or zoospores, and resting stages are not clearly known.

#### Excentrochloris

*Excentrochloris* is a solitary, often large, multinucleate coccoid, irregular, or ellipsoid unicellular genus. Adult cells may show local wall thickenings and partial protoplast segregation, but complete cellular division into an integrated multicellular thallus is not shown. Reproduction is by autospores and zoospores.

#### Pseudopleurochloris

*Pseudopleurochloris* is a coccoid unicellular xanthophycean genus described from Antarctic pack ice. It is treated as unicellular because its organization is based on individual coccoid cells rather than persistent filaments or coenocytic thalli. Reproductive interpretation follows related *Pleurochloris*-like genera, with autospore or zoospore reproduction reported.

#### Sphaerosorus

*Sphaerosorus* is interpreted as a spherical or grouped unicellular to colonial form. The genus is not treated as multicellular unless a species demonstrates persistent physical integration beyond mucilage association or parental-wall retention. Reproductive bodies are described as multiple autospores.

#### Mischococcus

*Mischococcus* is a unicellular or colonial, mucilage-associated genus in which the mucilaginous base functions primarily as an attachment structure. The occurrence of grouped cells does not by itself indicate true multicellularity, because the cells are not organized into a necessary serially integrated thallus. Reproduction is reported to involve zoospores and autospores.

#### Botryochloris

*Botryochloris* is treated as a coccoid or colonial form rather than a true multicellular filament. Older cells become multinucleate. Cell groups may be conspicuous, but the relevant organization is aggregation or retention of cells rather than developmentally necessary serial connection. Propagules are autospores or zoospores.

#### Pleurochloris

*Pleurochloris* is a coccoid unicellular xanthophycean genus. Cells may occur singly or in small groups, and older cells may become multinucleate, but this multinuclear condition is interpreted as a temporary cell cycle feature like *Botrydiopsis*, rather than multicellularity. Reproduction involves zoospores and autospores in larger cells.

#### Botrydiopsis

*Botrydiopsis* is a giant unicellular coccoid genus. Young cells may be uninucleate, and older cells may become strongly multinucleate with many chloroplasts, but this condition is not treated as multicellularity because the body remains a single cell with no cell division or functional differentiation. Reproduction may involve many zoospores or autospores.

#### Asterosiphon

*Asterosiphon* is a branched coenocytic alga with an integrated siphonous body plan and a rhizoid anchoring system. It is treated as multicellular-grade because it shows organism-level architecture with simple differentiation (i.e., akinetes formation and rhizoid) rather than a simple solitary multinucleate cell. Reproduction is mainly vegetative through differentiated propagule pathways, including akinete, aplanospore, and autospore stages reported in classical accounts. Flagellated zoospores have not been reported, with only non-flagellated amoeboid spores documented.

#### Botrydium

*Botrydium* is a coenocytic terrestrial or amphibious alga with an aerial vesicle and rhizoidal system. It is treated as multicellular-grade because the thallus is macroscopic, regionally differentiated, and developmentally integrated, even though regular septate cells are absent. Reproduction includes zoospores, aplanospores, and akinetes, including resistant stages in rhizoids.

#### Ophiocytium

*Ophiocytium* is generally a unicellular genus despite its remarkable cell elongation and occasional colony-like arrangements. The organism may become large, curved, stalked, or grouped, but the vegetative unit is a single cell rather than a serial multicellular filament. Old reports of multiple internal aplanospores should be interpreted cautiously, because multiple small non-motile bodies produced by internal cleavage as miniatures may correspond developmentally to autospores.

#### Xanthonema (syn. Heterothrix Pascher, 1932)

*Xanthonema* and historically related *Heterothrix*-like concepts represent filamentous multicellular organization because cells form persistent linear thalli. Zoospores are reported, and monospore-type aplanospores or akinetes occur in some species.

#### Bumilleria

*Bumilleria* forms short or long simple filaments and therefore represents a filamentous multicellular organization when persistent cell rows are present. Many species are fragile and readily fragment, so vegetative propagation by breakage happens. Zoospores are reported, and filament akinetes may occur in some taxa.

#### Bumilleriopsis

*Bumilleriopsis* is usually best described as unicellular to weakly colonial rather than clearly multicellular. Cells may be elongate and may remain temporarily aligned after autospore production, but such associations do not necessarily form a persistent developmental filament. However, *Bumilleriopsis filiformis* can be treated as multicellular because it frequently forms long filamentous cell chains produced. This creates a persistent serial thallus composed of multiple connected cells, unlike other *Bumilleriopsis* species that are mainly solitary unicells reproducing by autospores. Reproduction commonly involves autospores and zoospores.

#### Pseudobumilleriopsis

*Pseudobumilleriopsis* is treated here as unicellular or weakly colonial unless a species shows repeated persistent filaments. Internal non-motile daughter cells can remain associated after reproduction, but only temporarily. The principal propagules are interpreted as autospores, with zoospores also reported.

#### Tribonema

*Tribonema* is a filamentous multicellular genus in which vegetative growth produces persistent serial cells. Zoospores are commonly produced from vegetative cells, and the number per cell varies among species. Akinete formation is frequent in many taxa. Classical descriptions sometimes report monospore-type aplanospores, but when multiple non-motile daughter bodies are produced inside one cell (e.g., two daughter spores in *Tribonema aequale*), those structures may be better interpreted as autospores under the developmental definition used here.

#### Vaucheria

*Vaucheria* is the clearest coenocytic multicellular-grade xanthophycean lineage. The thallus is macroscopic, polarized, and differentiated, and it forms specialized reproductive organs such as antheridia and oogonia. Cytoplasmic and nuclear transport, localized septation, wound responses, synzoospore formation (e.g., in *Vaucheria bursata*), monospore-type aplanospores, and oogamous sexual reproduction all support interpretation as an integrated multicellular-grade siphonous organism.

